# Identification and structural characterization of a novel acetyl xylan esterase from *Aspergillus oryzae*

**DOI:** 10.1101/2024.08.23.609331

**Authors:** Chihaya Yamada, Tomoe Kato, Yoshihito Shiono, Takuya Koseki, Shinya Fushinobu

## Abstract

Acetyl xylan esterase plays a crucial role in the degradation of xylan, the major plant hemicellulose, by liberating acetic acid from the backbone polysaccharides. Acetyl xylan esterase B from *Aspergillus oryzae*, designated *Ao*AXEB, was biochemically and structurally investigated. The *Ao*AXEB-encoding gene with a native signal peptide was successfully expressed in *Pichia pastoris* as an active extracellular protein. The purified recombinant protein had pH and temperature optima of 8.0 and 30 °C, respectively, and was stable up to 35°C. The optimal substrate for hydrolysis by purified recombinant *Ao*AXEB among a panel of α-naphthyl esters was α-naphthyl acetate. Recombinant *Ao*AXEB catalyzes the release of acetic acid from wheat arabinoxylan. The release of acetic acid from wheat arabinoxylan increases synergistically with xylanase addition. No activity was detected using the methyl esters of ferulic, *p*-coumaric, caffeic, or sinapic acids. The crystal structures of *Ao*AXEB in the apo and succinate complexes were determined at resolutions of 1.75 and 1.90 Å, respectively. Although *Ao*AXEB has been classified in the Esterase_phb family in the ESTerases and alpha/beta-Hydrolase Enzymes and Relatives (ESTHER) database, its structural features partly resemble those of ferulic acid esterase in the FaeC family. Phylogenetic analysis also indicated that *Ao*AXEB is located between the clades of the two families. Docking analysis provided a plausible binding mode for xylotriose substrates acetylated at the 2- or 3-hydroxy position. This study expands the repertoire of side chain-degrading enzymes required for complete plant biomass degradation.

## Introduction

Xylans are the primary constituents of hemicelluloses and, after cellulose, are the second most abundant renewable polysaccharide in nature. Xylans conventionally contain heterogeneous substituents, such as arabinose, 2- and 3-*O*-acetyl groups, ferulic (4-hydroxy-3-methoxycinnamic), *p*-coumaric (4-hydroxycinnamic), and 4-*O*-methylglucuronic acids [1]. Acetyl xylan esterases (AXEs, EC 3.1.1.72) hydrolyze ester linkages to release acetic acid from acetylated xylans [2–4], whereas ferulic acid esterases (FAEs, EC 3.1.1.73) hydrolyze ester-linked ferulic acids. Most of the carboxylesterases adopt the α/β-hydrolase fold as a versatile scaffold and have a Ser-His-Asp (Glu) catalytic triad or a Ser-His catalytic dyad in their active sites for the catalytic mechanism using Ser as the nucleophile [5]. The ESTerases and alpha/beta-Hydrolase Enzymes and Relatives (ESTHER) database is dedicated to the α/β-hydrolase fold enzymes and currently classifies 248 subfamilies [6]. AXEs are classified into seven families (Abhydrolase_7, Acetyl-esterase_deacetylase, Acetylxylan_esterase, Antigen85c, Esterase_phb, Cutinase_like, and FaeC families) in the ESTHER database. AXEs belong to nine carbohydrate esterase (CE) families (CE1–CE7, CE12, and CE16) in the Carbohydrate-Active enZYmes (CAZy) database [7]. Most of the characterized enzymes belong to CE1 [8], and further subfamily classifications of fungal CE1 have been proposed [9]. According to the CAZy database classification, the AXEs from *Aspergillus* spp. belong to CE1 [10–13] and CE16 [14]. AXEs from *Aspergillus ficuum, Aspergillus luchuensis*, *Aspergillus oryzae, Podospora anserina*, and *Parastagonospora nodorum* heterologously expressed in *Pichia pastoris* have been characterized [9–12,15]. AXEs in CE2, CE3, CE6, CE12, and CE16 are also classified into the SGNH hydrolase subfamily of the GDSL family [16], which has a conserved GDSL motif around the nucleophilic serine residue instead of the canonical GxSxG motif present in other serine esterases [17]. Notably, as a fungal AXE, the crystal structure of *A. luchuensis* AXE A (*Al*AXEA) that is classified into Esterase_phb and CE1 families has been only reported [18], whereas several crystal structures of bacterial SGNH-type AXEs are solved to date.

In the present study, we identified a new esterase from *A. oryzae* that has low sequence similarity to characterized AXEs. Gene AO090005000945 encodes a hypothetical protein in the *A. oryzae* genomic database (http://www.bio.nite.go.jp/dogan/). Here, we report the identification and characterization of a novel AXE, designated *Ao*AXEB. We determined the crystal structure of *Ao*AXEB to investigate the structural basis of the classification and catalysis of this enzyme.

## Results and Discussion

### Expression of *AoaxeB* gene and purification of recombinant *Ao*AXEB

A comparison of the amino acid sequence of *A. oryzae* ORF AO090005000945, provisionally named *AoaxeB*, with the protein database revealed higher sequence identity to hypothetical proteins from *Aspergillus flavus*, *Aspergillus clavatus*, and *Aspergillus glaucus* (Table 1). Low sequence identity was observed with characterized AXEs or FAEs belonging to the FaeC and Esterase_phb families: *A. oryzae* AXE C (*Ao*AXEC) [15], *Talaromyces funiculosus* FAE A (*Tf*FAEA; accession number CAC85738.1), and *A. oryzae* FAE D (*Ao*FaeD) [19] in FaeC, and *A. oryzae* AXE A (*Ao*AXEA) [11] and *Al*AXEA [18] in Esterase_phb. Despite its low sequence similarity, AO090005000945 is currently listed as a putative FAE/poly(3-hydroxybutyrate) depolymerase/AXE in the Esterase_phb family (ID: aspor-q2ur69) in the ESTHER database. The CAZy database does not list this ORF within the CE families.

**Table 1.**
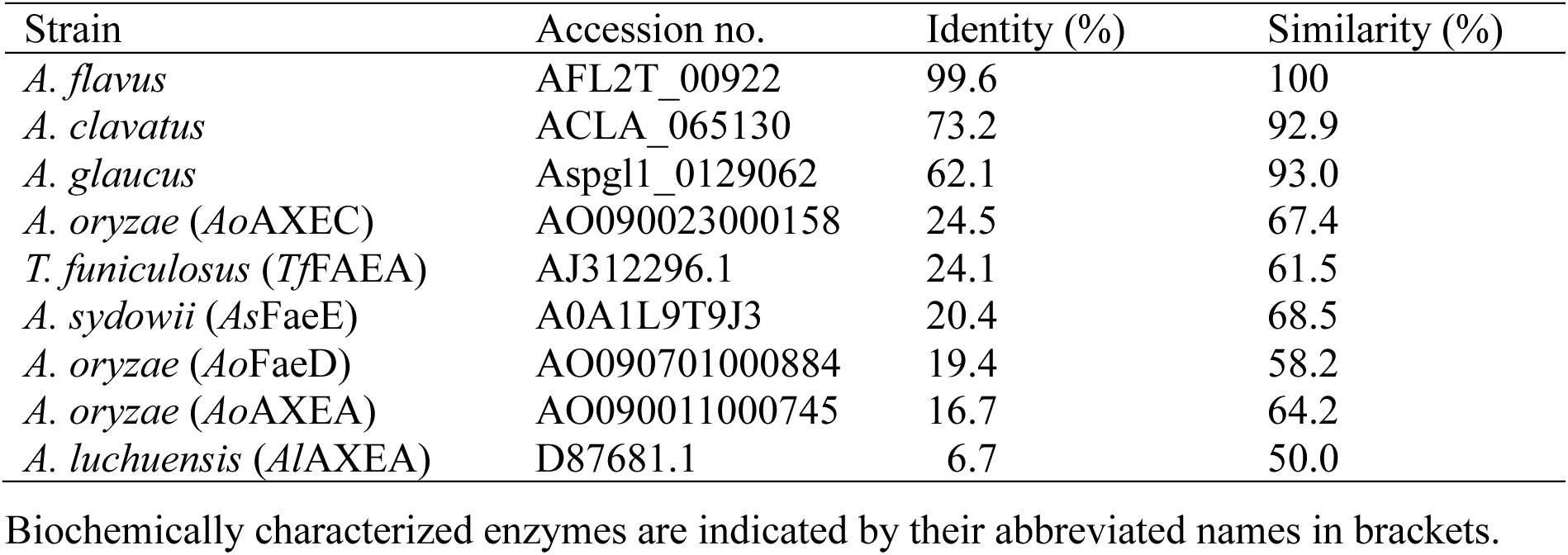
Fungal AXEs and FAEs showing similarity to *Ao*AXEB.

*A. oryzae* ORF AO090005000945, including the original signal sequence, was successfully engineered for protein expression in the heterologous host *P. pastoris.* The recombinant protein of *Ao*AXEB was secreted into the medium as an active enzyme and purified using a two-step procedure involving anion exchange and gel filtration chromatography. Purified *Ao*AXEB before and after treatment with endo-β-*N*-acetylglucosaminidase H (Endo-H) migrated using in dodecyl-sulfate polyacrylamide gel electrophoresis (SDS-PAGE) with a molecular mass of approximately 43 kDa and 34 kDa, respectively (Fig. 1), suggesting that the enzyme possessed *N*-linked oligosaccharides. The NetNglyc 1.0 server predicted two putative *N*-glycosylation sites (N42 and N48) in the *Ao*AXEB protein (Supplementary Table S1).

**Fig. 1.**
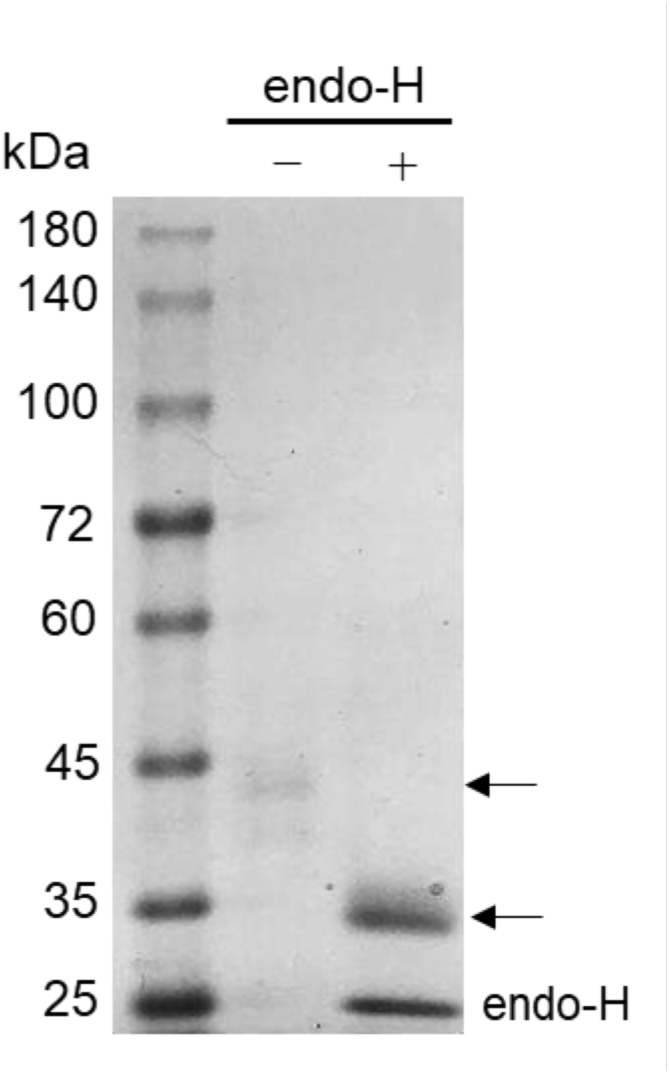
Coomassie-stained SDS-PAGE of purified *Ao*AXEB. Lane 1, molecular mass markers; lane 2, purified *Ao*AXEB; and lane 3, Endo-H-treated *Ao*AXEB.

### General properties of the purified *Ao*AXEB

The optimum pH of purified *Ao*AXEB was 8.0 with α-naphthyl acetate (C2) as the substrate (Fig. 2A), indicating that *Ao*AXEB had a higher optimum pH than *Ao*AXEA and *Ao*AXEC [12,15]. The optimum temperature for *Ao*AXEB activity was 30 °C. Thermal stability studies ranging 25–50 °C were performed in 50 mM sodium phosphate buffer (pH 8.0) (Fig. 2B). *Ao*AXEB was stable up to 35 °C. However, the thermal stability decreased to 50% of residual activity for 1 h incubation at 40 °C. The thermal stability of *Ao*AXEB was significantly lower than those of *Ao*AXEA and *Ao*AXEC [12,15].

**Fig. 2.**
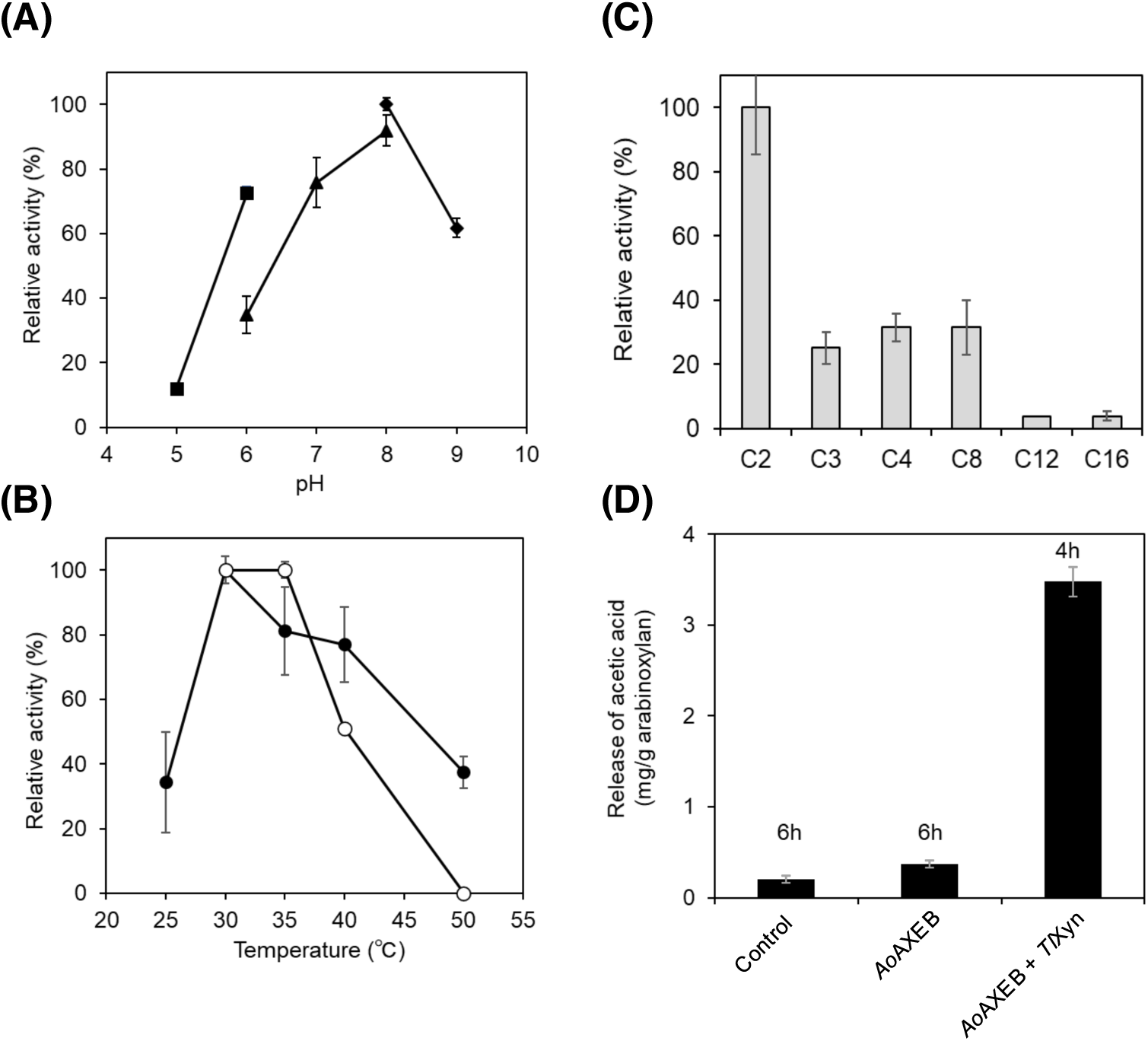
Biochemical properties of *Ao*AXEB. (A) Effects of pH on activity. Enzyme activity was measured in citrate buffer (pH 5–6) (closed squares), phosphate buffer (pH 6–8) (closed triangles), and Tris-HCl buffer (pH 8–9) (closed diamonds) at 30 °C using α-naphthyl acetate as a substrate. (B) Effects of temperature on activity (closed circles) and stability (open circles). Enzyme activity was measured in phosphate buffer (pH 8) at different temperatures. For the stability assay, aliquots of purified *Ao*AXEB were incubated for 1 h at different temperatures. After cooling on ice, residual enzyme activity was measured at pH 8.0. (C) Substrate specificity measured using different acyl chain α-naphthyl esters (C2–C16) as chromogenic substrates. Observed maximum activity was set at 100%. (D) Activity toward acetylated xylan. Release of acetic acid from insoluble wheat arabinoxylan was determined as described in the Materials and Methods section.

### Substrate specificity

The hydrolytic activity of the purified recombinant *Ao*AXEB was examined using a panel of α-naphthyl esters (C2–C16) as artificial substrates. The optimal substrate for *Ao*AXEB was α-naphthyl acetate (C2), and it showed lower activity toward acyl chain substrates containing three or more carbon atoms (Fig. 2C). The specific activity of *Ao*AXEB toward the C2 substrate (0.033 ± 0.005 units/mg protein) was three-fold higher than that toward α-naphthyl butyrate (C4, 0.011 ± 0.001 units/mg protein). This result suggests that *Ao*AXEB shows a higher specificity for acetic acid esters than *Ao*AXEC [15], although the specific activity of *Ao*AXEB is lower than that of *Ao*AXEC. The *K*_m_ and *k*_cat_ values toward the C2 substrate were 0.24 ± 0.12 mM and 0.17 ± 0.03 s^-1^ for *Ao*AXEB and 1.9 ± 0.4 mM and 4.5 ± 0.7 s^-1^ for *Ao*AXEC, respectively. Therefore, the catalytic efficiency (*k*_cat_/*K*_m_) for *Ao*AXEB (0.72 s^-1^°mM^-1^) was lower than that for *Ao*AXEC (2.4 s^-1^°mM^-1^). Methyl ferulate, methyl *p-* coumarate, methyl caffeate, and methyl sinapate were not hydrolyzed, indicating that *Ao*AXEB did not exhibit FAE activity.

### Release of acetic acid from acetylated xylan and synergism with xylanase

When *Ao*AXEB was incubated with wheat arabinoxylan at 37 °C for 6 h, the amount of released acetic acid (0.37 ± 0.04 mg/g substrate) increased by 1.85-fold compared to the enzyme-free condition (0.20 ± 0.04 mg/g) (Fig. 2D). A combination of *Ao*AXEB and *Thermomyces lanuginosus* xylanase (*Tl*Xyn) considerably released acetic acid (3.48 ± 0.16 mg/g substrate) at 37 °C for 4 h. Thus, the amount of acetic acid released during a shorter incubation with the xylanase was 9.5-fold the amount released during the incubation with *Ao*AXEB alone. These observations suggest that *Ao*AXEB deacetylates xylooligomers instead of xylan polymers. In contrast, *Ao*AXEC acts directly on the acetylated xylan polymer [15]. However, no significant synergistic effect of *Ao*AXEB was observed in the degradation of wheat arabinoxylan by *Tl*Xyn (Supplementary Fig. S1). In contrast, the release of acetic acid from xylan increased synergistically upon the addition of xylanase to *Ao*AXEA [12].

### Crystal structure

The crystal structures of recombinant *Ao*AXEB were determined in apo and succinate complex forms at 1.75 and 1.90 Å resolutions, respectively (Table 2). The crystals of both structures contained one molecule of *Ao*AXEB in the asymmetric unit. The PISA server [20] predicted that the biological assembly in the solution was a monomer. Fig. 3A shows the overall structure of the succinate complex form. A clear electron density map of the succinate molecule was observed at the active site (Fig. 3B), where the apo structure contained water molecules (Supplementary Fig. S2). The succinate molecule may have been derived from trace contamination in the acetate reagent used for crystallography or the fermentation product of *P. pastoris* used for recombinant protein production because we did not use succinate in the crystallization solution. An acetate molecule was observed on the protein surface because sodium acetate buffer (100 mM) was used in the crystallization solution (Fig. 3A). The apo- and succinate-complex structures are similar, although there is a structural difference far from the active site (Supplementary Figs. S3A and B). The root mean square deviations (RMSD) between the two structures was 0.13 Å for 253 over 298 Cα atoms, and there was no structural difference near the active site. The structure predicted using AlphaFold was closer to the apo form (Supplementary Fig. S3C).

**Fig. 3.**
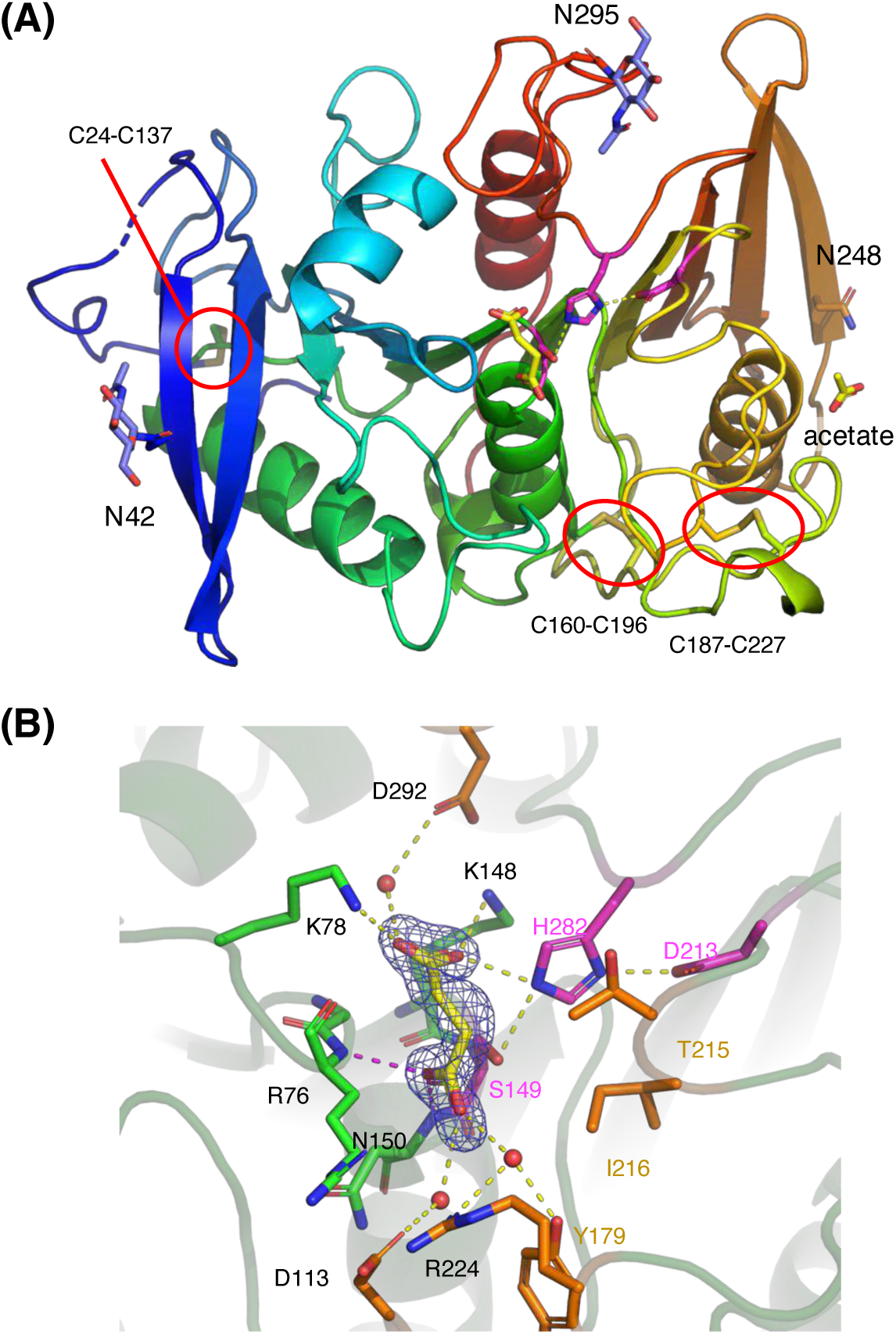
Crystal structure of *Ao*AXEB. (A) Overall structure. Polypeptides are shown in rainbow colors. *N*-Glycans (blue), catalytic triad (magenta), disulfide bonds (sulfur atoms in yellow, indicated with red circles), succinate (yellow), and acetate (yellow) are shown as sticks. (B) Active site. A Polder map (4σ) is shown for the bound succinate. Water atoms and hydrogen bonds are shown as red spheres and yellow dotted lines, respectively. Residues forming direct hydrogen bonds with succinate and other active site residues are shown as green and orange sticks, respectively.

**Table 2.** Crystallographic data collection and refinement statistics.

|  | *AoAXEB apo* | *AoAXEB + succinate* |
| --- | --- | --- |
| Data collection* |  |  |
| Beamline | KEK PF BL5A | SLS PSI X06DA |
| Wavelength (Å) | 1.000 | 1.000 |
| Space group | $P2_12_12_1$ | $P2_12_12_1$ |
| Unit cell (Å) | $a = 57.614, b = 63.529, c = 83.417$ | $a = 57.027, b = 64.008, c = 83.324$ |
| Resolution (Å) | 47.41–1.75 | 47.06–1.90 |
| Total reflections | 207,748 (10,264) | 133,539 (2,020) |
| Unique reflections | 31,605 (1,710) | 23,666 (1,011) |
| $R_{\text{merge}}$ | 0.117 (0.859) | 0.086 (0.219) |
| $R_{\text{pim}}$ | 0.049 (0.379) | 0.037 (0.176) |
| Mean $I/\sigma(I)$ | 12.1 (2.1) | 13.7 (4.1) |
| $CC_{1/2}$ | 0.998 (0.713) | 0.994 (0.715) |
| Completeness (%) | 100.0 (100.0) | 96.0 (65.8) |
| Multiplicity | 6.6 (6.0) | 5.6 (2.0) |
| Wilson B-factor (Å<sup>2</sup>) | 9.60 | 11.56 |
| Refinement |  |  |
| Resolution (Å) | 45.45–1.75 | 47.10–1.90 |
| Reflections | 31,549 | 23,621 |
| $R_{\text{work}}/R_{\text{free}}$ | 0.164/0.199 | 0.165/0.199 |
| Number of atoms |  |  |
| Amino acids | 2,318 | 2,307 |
| Ions | 1 (K<sup>+</sup>) | 0 |
| Ligands | 52 (1 PGE, 3 NAG) | 40 (1ACT, 1SIN, 2NAG) |
| Waters | 323 | 245 |
| Glycosylation | N42, N248, N295 | N42, N295 |
| Average B-factor (Å<sup>2</sup>) |  |  |
| Protein | 17.28 | 16.3 |
| Ligands | 37.25 | 38.23 |
| Ions | 23.5 | – |
| Waters | 26.1 | 23.08 |
| Clashscore | 1.30 | 2.86 |
| Molprobity score | 0.99 | 1.39 |
| Ramachandran plot (%) |  |  |
| Favored | 97.98 | 95.92 |
| Allowed | 1.68 | 3.06 |
| Outlier | 0.34 | 1.02 |
| RMSD from ideal values |  |  |
| Bond lengths (Å) | 0.0149 | 0.0103 |
| Bond angles (°) | 1.84 | 1.62 |
| PDB code | 9J07 | 9J08 |

\*Values in parentheses represent the highest resolution shell.

*Ao*AXEB consists of a single domain adopting a typical α/β-hydrolase fold, in which a 10-stranded β-sheet is sandwiched by several helices (Fig. 3A). There were three disulfide bonds (C24-C137, C160-C196, and C187-C227). Because we used a protein sample treated with Endo-H, a single *N*-acetylglucosamine was observed at the *N*-glycosylation sites. The apo-structure was *N*-glycosylated at N42, N248, and N295 (Supplementary Fig. S3A), whereas the succinate complex contains two *N*-linked glycans at N42 and N295 (Fig. 3A and Supplementary Fig. S3B). This observation partly contradicts the prediction by NetNglyc (Supplementary Table S1), where N42 and N248 are likely *N*-glycosylation sites, and N295 and N321 are not.

The catalytic triad is located at the center of the molecule and consists of S149, H282, and D213 (Fig. 3B). The succinate molecule was bound to the active site through numerous hydrophilic interactions. One carboxy group forms direct hydrogen bonds with the side chains of K78, K148, and H282 and a water-mediated hydrogen bond with D292. The other carboxy group forms direct hydrogen bonds with the main chain amides of R76 and N150 and water-mediated hydrogen bonds with D113, Y179, and R224. The interaction with the main chain amides of R76 and N150 corresponds to an oxyanion hole, which plays a key role in the catalysis of α/β-hydrolases (discussed below).

### Structural comparison

A Dali structural similarity search revealed that *Ao*AXEB was most similar to *Lihuaxuella thermophila* poly[(*R*)-3-hydroxybutyric acid] depolymerase (*Lt*PHBase), which belongs to the Esterase_phb family (Table 3) [21]. The second hit was *Acremonium alcalophilum* FAE D (*Aa*FaeD), which belongs to the FaeC family [22]. The third and fourth hits were *Aspergillus sydowii* FAE E (*As*FaeE) [23] and *Al*AXEA, respectively [18], both of which belong to Esterase_phb. The fifth hit was a thermostable esterase from *Thermotoga maritima* (*Tm*EstA) belonging to the 5_AlphaBeta_hydrolase family [24], which showed significantly lower structural similarity than the four proteins listed above. The overall structures of *Ao*AXEB and its five structural homologs are shown in Fig. 4. Interestingly, two of the three disulfide bonds in *Ao*AXEB (C24-C137 and C160-C196) were conserved with *Aa*FaeD in the FaeC family (Figs. 4A and C). Esterase_phb proteins have a conserved disulfide bond near the catalytic site (C40-C75 in *Al*AXEA, Figs. 4B, D, and E) [18], which is not present in *Ao*AXEB. A disulfide bond corresponding to C160-C196 in *Ao*AXEB was also found in *As*FaeE and *Al*AXEA but not in the most structurally similar *Lt*PHBase. In summary, *Ao*AXEB has structural features that are intermediate between those of the Esterase_phb and FaeC families. Phylogenetic tree analysis of the amino acid sequences also indicated that *Ao*AXEB was located between two families (Fig. 5).

**Fig. 4.**
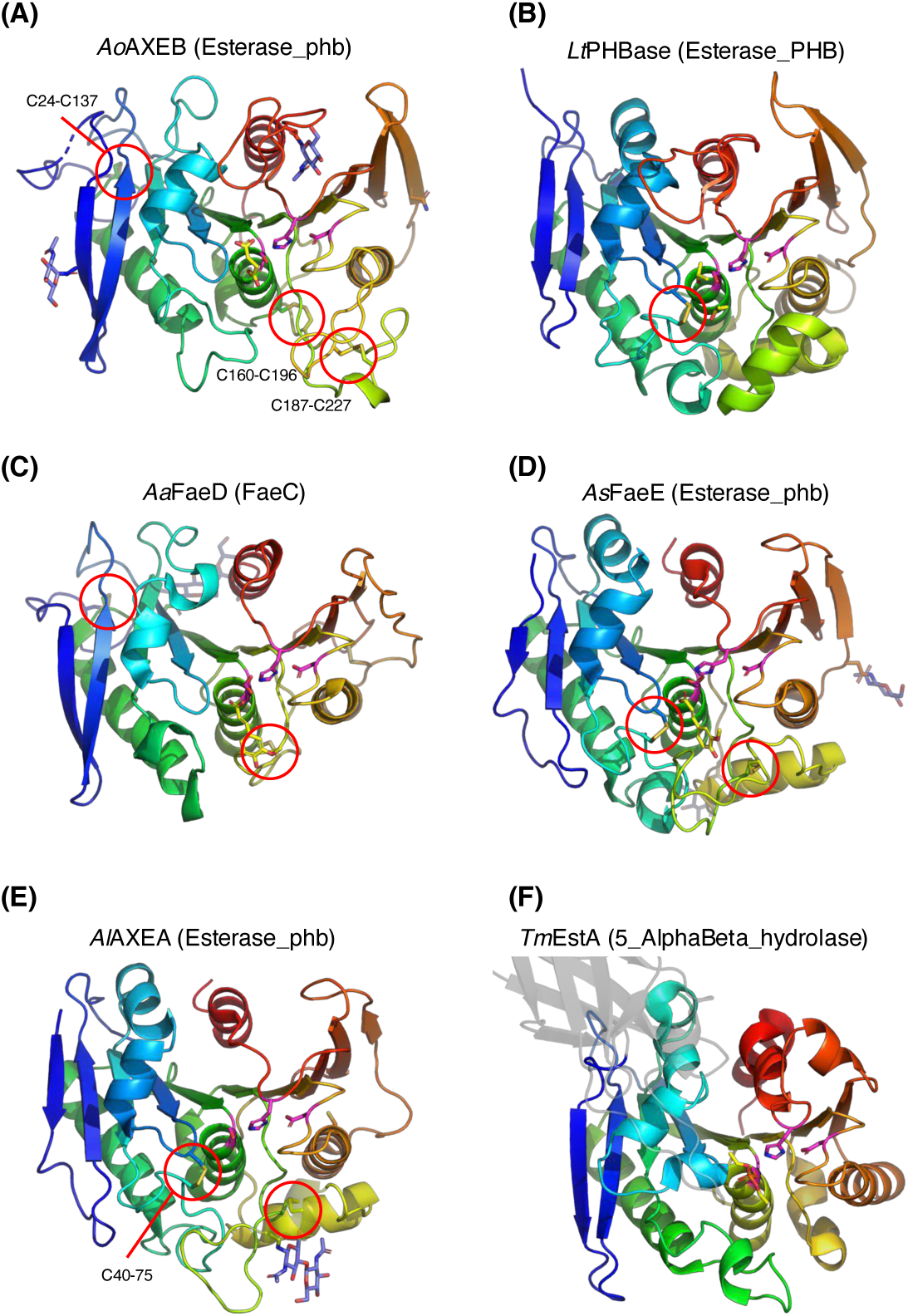
Structural comparison with other α/β-hydrolase family enzymes in the top five hits of the structural similarity search. (A) *Ao*AXEB complexed with succinate. (B) *L. thermophila* poly[(*R*)-3-hydroxybutyric acid] depolymerase (*Lt*PHBase) complexed with isopropanol (PDB ID: 8DAJ). (C) *A. alcalophilum* FAE D (*Aa*FaeD) complexed with ferulic acid (PDB ID: 8JH9). (D) *A. sydowii* FAE E (*As*FaeE) complexed with ferulic acid (PDB ID: 8IYB). (E) *A. luchuensis* AXE A (*Al*AXEA, PDB ID: 5X6S). (F) *T. maritima* esterase (*Tm*EstA) complexed with diethyl phosphate (PDB ID: 3DOI). The N-terminal Ig-like domain is shown transparently in grey. *N*-Glycans (blue), catalytic triad (magenta), disulfide bonds (sulfur atoms in yellow and marked with red circles), and bound ligands (yellow) are shown as sticks.

**Fig. 5.**
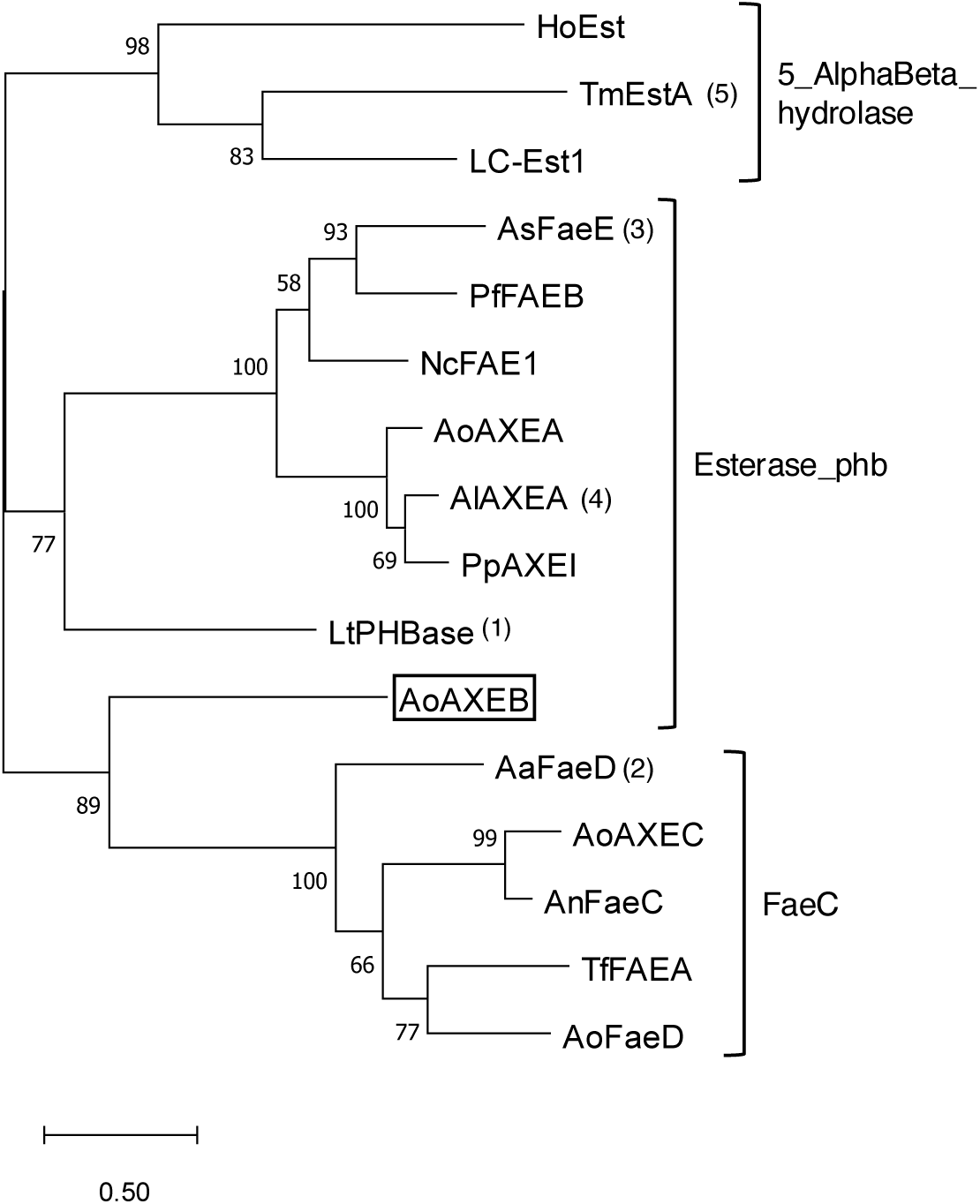
A phylogenetic tree of esterases in Esterase_phb, FaeC, and 5_AlphaBeta_hydrolase families. Order of structural similarity with *Ao*AXEB is indicated in parentheses. A maximum likelihood tree with the highest log likelihood (-10132.60) is shown. This analysis involved 16 amino acid sequences, with a total of 581 positions in the final dataset. Bootstrap values of 200 replications are indicated at the branches. The scale indicates branch lengths measures in the number of substitutions per site. The protein names not listed in the text and their ESTHER database IDs are as follows: HoEST, *Haliangium ochraceum* phospholipase/carboxylesterase (halo1-d0lmj0); LC-Est1, a metagenome-derived esterase (9bact-3WYDseq) [38]; PfFAEB, *Penicillium funiculosum* FAE B (penfn-faeb) [39]; NcFAE1, *Neurospora crassa* Fae-1 (neucr-faeb) [40]; PpAXEI, *Penicillium purpurogenum* AXE I (penpu-AXEI) [41]; and AnFaeC, *Aspergillus nidulans* putative FAE (emeni-faec).

**Table 3.** Result of DALI structural similarity search.

| Protein | ESTHER<sup>a</sup> | CAZy<sup>b</sup> | PDB ID<br>(chain) | Z score | RMSD<br>(Å) | LALI<sup>c</sup> | %ID<sup>d</sup> |
| --- | --- | --- | --- | --- | --- | --- | --- |
| *LtPHBase* | Esterase_phb | NL | 8daj (A) | 28.8 | 2.4 | 253 | 21 |
| *AaFaeD* | FaeC | NL | 8jh9 (A) | 27.9 | 2.3 | 234 | 26 |
| *AsFaeE* | Esterase_phb | CE1 | 8iyb (A) | 24.4 | 2.3 | 226 | 21 |
| *AlAXEA* | Esterase_phb | CE1 | 5x6s (B) | 23.8 | 2.5 | 229 | 20 |
| *TmEstA* | 5_AlphaBeta_<br>hydrolase | NL | 3doh (A) | 20.8 | 2.2 | 206 | 18 |

2 Analyzed using the DALI server (http://ekhidna2.biocenter.helsinki.fi/dali/).
<sup>a</sup>Rank 1 family in the ESTHER database (https://bioweb.supagro.inrae.fr/ESTHER/). <sup>b</sup>Family in the CAZy database (http://www.cazy.org). NL, not listed;. <sup>c</sup>Number of aligned residues.
<sup>d</sup>Sequence identity.

### Active site features

The active sites of *Ao*AXEB were compared with those of FAEs and AXE (Figs. 6A–D). The structures of AaFaeD and AsFaeE complexed with ferulic acid showed an open pocket or surrounding pocket for large aromatic compounds (Figs. 6B and C). In contrast, the putative active site of *Ao*AXEB had a small pocket surrounded by R76, R224, Y179, I216, and T215 (Fig. 6A). The pocket size of *Ao*AXEB is similar to that of *Al*AXEA (Fig. 6D). Our previous study showed that the small pocket of *Al*AXEA surrounded by the C40-C75 disulfide bond and the large aromatic side chain of W160 accommodated an acetyl group, because site-directed mutants at W160 acquired FAE activity [18].

**Fig. 6.**
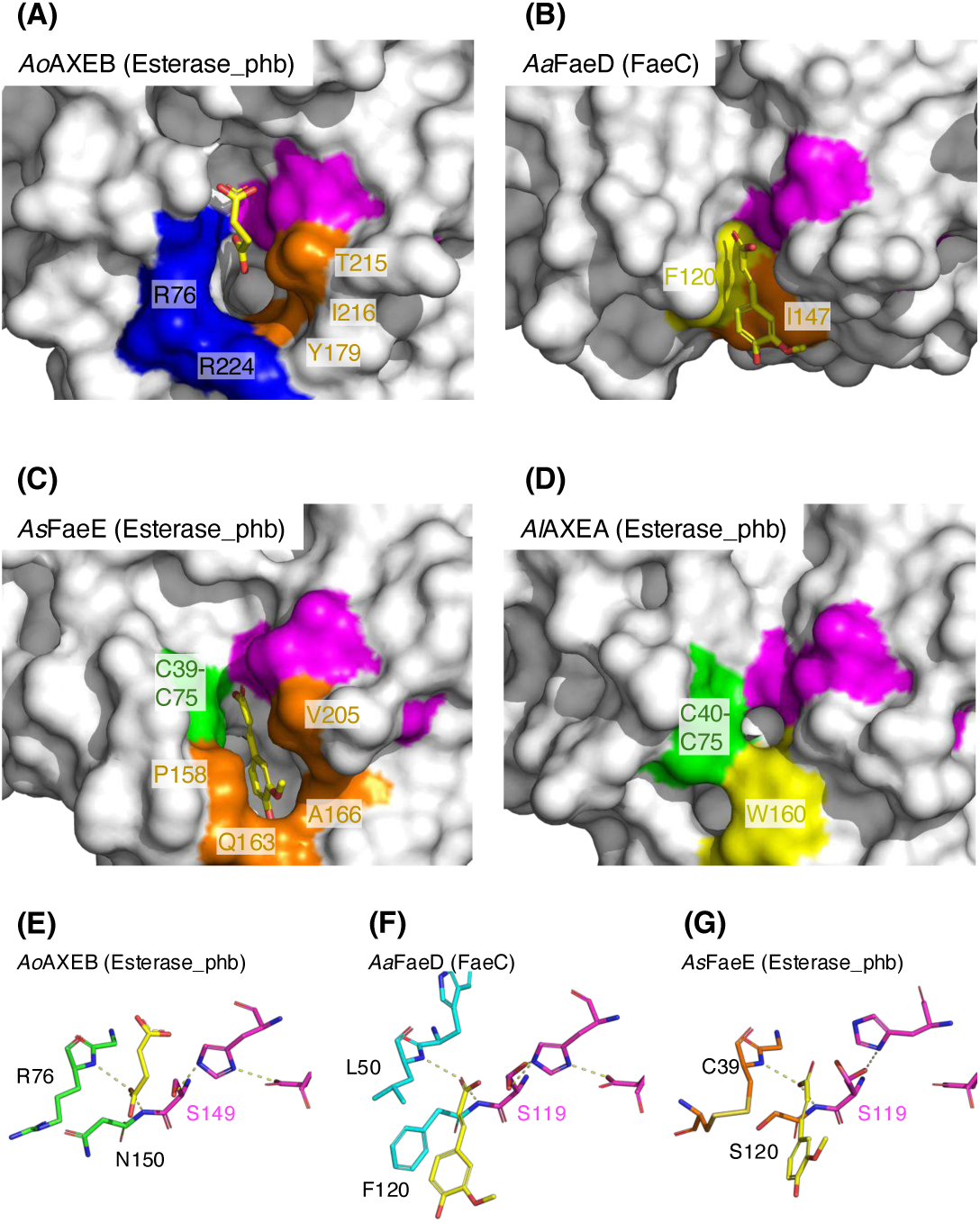
Active site comparison with FAEs and an AXE. (A–D) Molecular surface presentations. (A) *Ao*AXEB complexed with succinate. (B) *Aa*FaeD complexed with ferulic acid (PDB ID: 8JH9). (C) *As*FaeE complexed with ferulic acid (PDB ID: 8IYB). (D) *Al*AXEA (PDB ID: 5X6S). The catalytic triad residues are indicated with magenta, and other residues forming the active site pocket are indicated with blue (basic residues), yellow (aromatic residues), green (cysteine residues forming a disulfide bond), or orange (other types of residues). (E–G). The catalytic components of *Ao*AXEB (E, green), *Aa*FaeD (F, cyan), and *As*FaeE (G, orange). Bound ligands (succinate or ferulic acid in yellow), the catalytic triad (magenta), and oxyanion hole are shown as sticks. Note that the catalytic serine in *Aa*FaeD and AsFaeE takes two alternative conformations, and the catalytic histidine in *As*FaeE takes a deviated conformation without forming a hydrogen bond with aspartate.

The oxyanion hole is crucial for the catalytic function of serine and cysteine hydrolases and plays a pivotal role in stabilizing the tetrahedral intermediate during the reaction, typically through hydrogen bonding with the backbone amides of the polypeptides [25]. The oxyanion hole of *Ao*AXEB is formed by the main chain amides of R76 and N150 (Fig. 6E). This is conserved between *Aa*FaeD and *As*FaeE (Figs. 6F and G) as hydrogen bond interactions with a carboxylate group in the ligands are similarly formed.

We performed automated docking analysis to elucidate the substrate-binding mode of *Ao*AXEB. AXEs liberate acetic acid from their esters with 2- or 3-hydroxy groups on the xylan backbone [4]. The deacetylase activity of *Ao*AXEB on acetylated xylooligosaccharides in wheat arabinoxylan is shown (Fig. 2D). Therefore, for the docking analysis, we used xylotriose models acetylated at the 2- or 3-position of the central sugar unit. As a result, plausible binding modes of the 2-acetylated xylotriose (2AcX_3_) and 3-acetylated xylotriose (3AcX_3_) were obtained (Fig. 7). The direction of the non-reducing to reducing ends of xylotriose was reversed in 2AcX_3_ and 3AcX_3_; however, the hydrogen bond interactions with K78, D292, K148, S149, T215, and R223 were conserved. Both the 4-hydroxy group at the non-reducing end and the 1-hydroxy group at the reducing end of 2AcX_3_ and 3AcX_3_ pointed toward the solvent, suggesting that *Ao*AXEB can bind xylooligosaccharides for longer than four sugar units. The acetyl group in the central sugar units of 2AcX_3_ and 3AcX_3_ penetrated the small pocket of the protein, occupying a suitable position under nucleophilic attack by S149 and stabilizing the tetrahedral intermediate by the oxyanion hole. The pocket appears to be able to accommodate an alkyl group with several carbon atoms, but is small enough to favor an acetyl group. Thus, the docking results explained the substrate specificity of *Ao*AXEB (Fig. 2C).

**Fig. 7.**
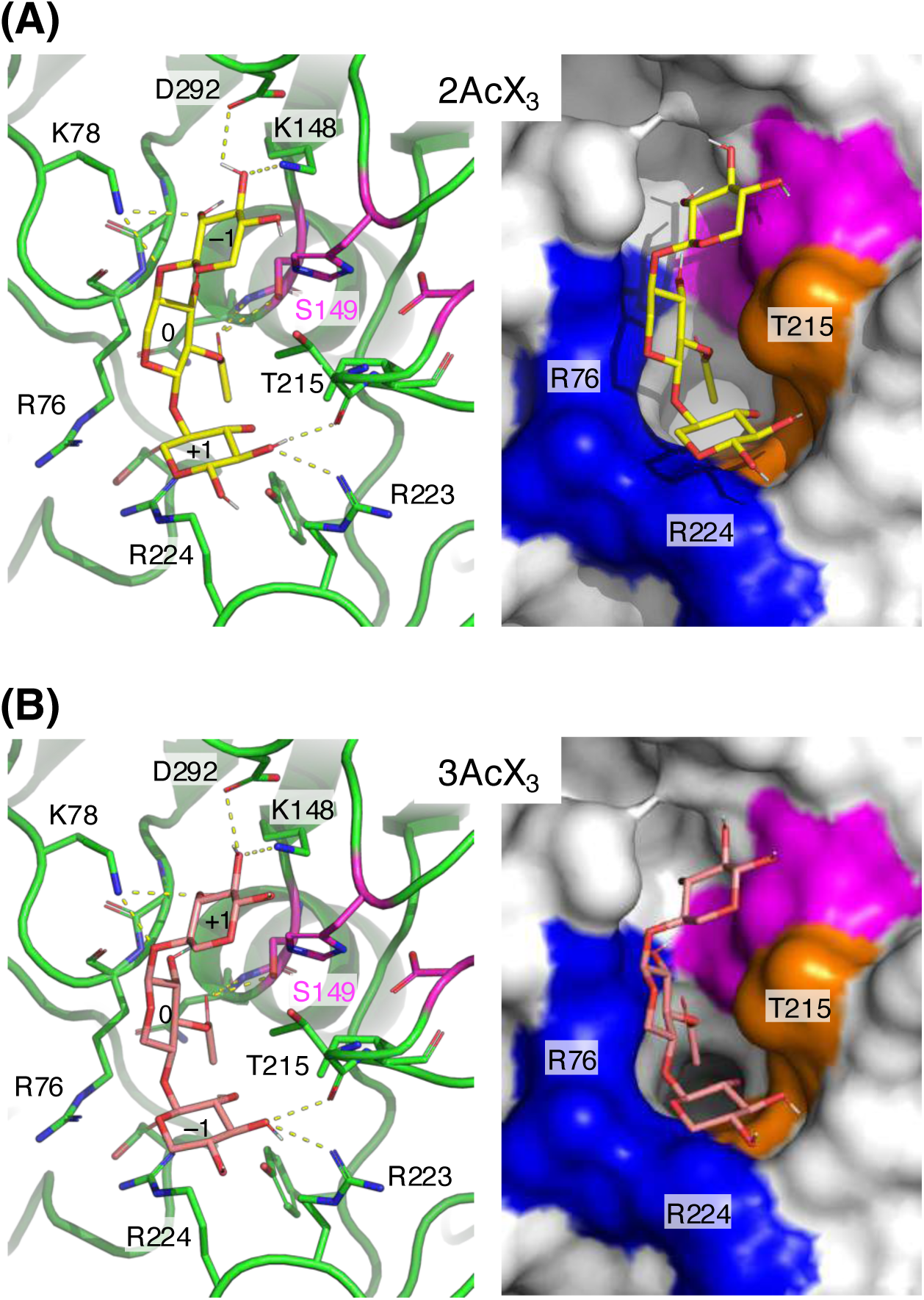
Possible binding modes of acetylated xylooligosaccharides in the active site of *Ao*AXEB. Results of automated docking analysis are shown as the molecular model (left) and surface (right). (A) 2-Acetylated xylotriose (2AcX_3_, yellow sticks). (B) 3-Acetylated xylotriose (3AcX_3_, pink sticks). Sugar units from the non-reducing to reducing ends are labeled with –1, 0, and +1. The middle sugar unit labeled “0” is acetylated.

## Conclusion

In this study, we discovered and characterized a novel AXE, *Ao*AXEB, from one of the most important industrial microorganisms, *A. oryzae* [26]. Although it has been listed as a putative FAE belonging to Esterase_phb in the ESTHER database, *Ao*AXEB exhibited typical substrate specificity for AXE activity and synergy with xylanase in the deacetylation of wheat arabinoxylan. The crystal structure of *Ao*AXEB revealed that it partly had structural features similar to those of FaeC, and phylogenetic analysis indicated that it was situated between the Esterase_phb and FaeC families. Docking analysis elucidated the possible binding mode of acetylated xylooligosaccharides at the active site. Our study suggests that there are still unexplored genes for plant cell wall-degrading enzymes in fungal genomes.

## Methods

### Strains and culture conditions

*A. oryzae* strain RIB40 and *P. pastoris* strain KM71H were used for the source of the gene cloning and the heterologous expression of the gene, respectively. *P. pastoris* transformants were grown at 30 °C in 10 mL of BMGY [1% yeast extract, 2% peptone, 1%(v/v) glycerol, 0.00004% biotin, 1.34% yeast nitrogen base with ammonium sulfate, and 10%(v/v) 1M potassium phosphate (pH 6.0)] medium in a shaking incubator until the cell density reached an OD_600_ of 4. Cells were harvested aseptically by centrifugation (3000×g, 10 min, 4 °C). The cells were then resuspended in 100 mL of BMMY medium (same composition as BMGY medium, except containing 0.5% (v/v) methanol instead of glycerol) in a 500 mL flask to an OD_600_ of 1 to start induction. The culture was kept in a shaking incubator at 30 °C for 7 d (180 rpm) with the addition of methanol (0.5 mL) once daily to maintain induction.

### Cloning and expression of *AoaxeB* gene

*AoaxeB* was annotated as the hypothetical protein AO090005000945 in the *A. oryzae* genomic database. The protein-coding sequence of *AoaxeB* was amplified using polymerase chain reaction (PCR) with the forward primer 5’-AG<u>GAATTC</u>ATGAAGTTTCTCTCAGTAAT-3’ (the *Eco*RI site is underlined), the reverse primer 5’-GT<u>TCTAGA</u>CTATCTCGCCTCGCTCTGGT-3’ (the *Xba*I site is underlined), and *A. oryzae* genomic DNA as a template. The PCR products digested with *Eco*RI and *Xba*I were cloned into an *Eco*RI–*Xba*I-digested pPICZB expression vector (Invitrogen, Waltham, MA, USA). The resulting construct pPICZB-AXEB was used to transform *Escherichia coli* DH5α, and the positive clone was confirmed using colony PCR. To splice an intron contained within the gene, PCR was performed by inverse PCR using forward primer 5’-AAAGAATGGCAAGGAGACCCA-3’ and reverse primer 5’-GTTGAGCCCCTGCGGGTACAC-3’ with the KOD-Plus-Mutagenesis Kit (TOYOBO, Otsu, Japan). Gene splicing was verified using DNA sequencing. This procedure yielded an expression plasmid vector containing the *AoaxeB* gene with its native signal sequence under the control of the alcohol oxidase 1 (*aox1*) promoter and terminator for expression in *P. pastoris* as described above.

### Purification of recombinant *Ao*AXEB

The recombinant plasmid (pPICZB-AXEB) was linearized with *Pme*I and subsequently transformed into *P. pastoris* KH71H using the Pichia EasyComp Transformation Kit (Invitrogen) according to the manufacturer’s protocol. The 7-d cell culture was harvested by centrifugation (5000×*g*, 15 min). The culture supernatant was used for purification of recombinant *Ao*AXEB. Enzyme purification was performed as previously described [15]. Enzyme purity and molecular mass were evaluated using 12% SDS-PAGE, followed by Coomassie Brilliant Blue staining. The protein concentration was measured using a Micro BCA Protein Assay Kit (Thermo Fisher Scientific Inc., USA). After boiling, the purified *Ao*AXEB protein was treated with 0.5 mU of Endo-H (FUJIFILM Wako Pure Chemical Industries, Osaka, Japan) in 50 mM sodium acetate buffer (pH 5.0) at 37 °C for 18 h.

### Assay of enzyme activity

The activity of purified *Ao*AXEB toward wheat arabinoxylan, α-naphthyl acetate (C2), propionate (C3), butyrate (C4), caprylate (C8), laurate (C12), palmitate (C16), methyl ferulate, methyl *p-*coumarate, methyl caffeate, and methyl sinapate was investigated as described previously [19,27]. The activity as a function of pH for the α-naphthyl acetate substrate was measured using citrate buffer (pH 5.0–6.0), phosphate buffer (pH 6.0–8.0), and Tris-HCl buffer (pH 8.0–9.0) at 30 °C. The activity as a function of temperature in the range 30–55 °C, using increments of 5.0 °C, was performed in 50 mM sodium phosphate (pH 8.0). To measure the thermal stability of *Ao*AXEB, the enzyme was incubated at an appropriate temperature for 1 h, and residual activity was determined using α-naphthyl acetate as the substrate.

### Release of acetic acid from acetylated xylan and synergism with xylanase

The activity of *Ao*AXEB toward acetylated xylan was assayed using insoluble wheat arabinoxylan (Megazyme, Braw, Wicklow, Ireland), at a final concentration of 2.0%, and in a final volume of 4.0 mL at 37 °C for 6 h in 50mM phosphate buffer (pH 8.0) with purified protein (60 μg). The amount of acetic acid released was determined using an F-kit acetate kit (Roche, Basel, Switzerland). The synergistic effect between AoAXEB and *Tl*Xyn (Sigma-Aldrich, St. Louis, MO, USA) was investigated by incubation of a 4 mL solution of 2.0% insoluble wheat arabinoxylan (Sigma-Aldrich) in 50 mM phosphate buffer (pH 8.0) for 4 h at 37 °C with purified *Ao*AXEB (60 μg) and *Tl*Xyn (10 μg). The synergistic effect was determined based on the amount of acetic acid released using an acetic acid assay kit (Roche).

### Crystallization and structure determination

The crystals of *Ao*AXEB were obtained at 20 °C using the sitting-drop vapor diffusion method by mixing equal volumes of protein and reservoir solutions. For apo-form crystals, a protein solution containing 2 mg/mL *Ao*AXEB and 10 mM xylooligosaccharides (FUJIFILM Wako Pure Chemical Industries) and a reservoir solution containing 0.2 M potassium fluoride and 20% PEG3350 (w/v) were used. The electron densities of the xylooligosaccharides were not observed in the resulting crystal structure. For succinate complex crystals, protein solution containing 3 mg/mL *Ao*AXEB and reservoir solution containing 0.1 M sodium acetate buffer (pH 4.5) and 20% (w/v) PEG3000 were used. For data collection, the crystals were cryoprotected using a reservoir solution supplemented with 20% (v/v) PEG200 and flash-cooled by dipping in liquid nitrogen. X-ray diffraction data were collected at 100 K on the beamlines at the Photon Factory of the High Energy Accelerator Research Organization (KEK, Tsukuba, Japan) and the Swiss Light Source (SLS) of the Paul Scherrer Institut (PSI) (Villigen, Switzerland). The preliminary diffraction data were collected at SPring-8 (Hyogo, Japan). The datasets were processed using the XDS [28] and Aimless software [29]. The initial phase was obtained through molecular replacement using MORDA [30] with the LC-Est1 structure (PDB ID: 3WYD, chain A) as a template. Phase improvement and automated model building were achieved using PHENIX (phase and build) [31] and BUCCANEER software [32]. Manual model rebuilding and refinement were performed using Coot (Emsley et al., 2010) and Refmac5 [33]. Polder maps were prepared using PHENIX software [34]. Molecular graphic images were prepared using PyMOL (Schrödinger LLC, New York, NY, USA).

### Phylogenetic and docking analyses

Phylogenetic analysis was performed using MEGA 11.0.13 [35]. The protein sequences were aligned using MUSCLE [36]. The docking study was performed using AutoDock Vina 1.2.5 [37]. Using AutoDockTools, polar hydrogen atoms were added to the amino acid residues, and Gasteiger charges were assigned to all atoms of the enzyme. The grid map was prepared with 20 × 20 × 20 points spaced at 1.0 Å distances. The grid box was centered on the carboxy carbon atom of succinate near the catalytic serine of *Ao*AXEB. The exhaustiveness value is 256. The ligand structure was docked at flexible torsion angles, whereas the protein structure was fixed. Docking of 2AcX_3_ yielded nine binding modes with estimated affinities ranging -6.958–(-6.336) kcal/mol, with the second-best result (-6.893 kcal/mol) being selected. Docking of 3AcX_3_ yielded eight binding modes with estimated affinities ranging -7.299–(-6.338) kcal/mol, with the second-best result (-6.893 kcal/mol) being selected. The first-ranked docking result was not catalytically competent for either ligand.

## Acknowledgments

The authors thank Dr. Takatoshi Arkakawa and Dr. Arnaud Chatonnet for their valuable discussions. We also thank the staff of KEK-PF, SLS at PSI, and SPring-8 for the X-ray data collection. This research was in part supported by the YU-COE program of Yamagata University (to TKo), JSPS-KAKENHI (19H00929 and 23H00322 to SF and 21K15025 to CY) and the Research Support Project for Life Science and Drug Discovery (Basis for Supporting Innovative Drug Discovery and Life Science Research (BINDS)) from AMED under Grant Number JP22ama121001.

## Author contributions

TKo, SF, and YS conceived and supervised the study; CY, TKo, and SF planned experiments; CY, TKa, and TKo performed experiments; TKa and TKo performed the protein production, purification, and biochemical experiments; CY and SF performed protein crystallography; SF performed the phylogenetic and docking analyses; and TKo and SF wrote the manuscript. All authors reviewed the final version of the manuscript.

## Data availability statement

Atomic coordinates and structure factors of the crystal structures have been deposited in the Protein Data Bank under accession numbers 9J07 and 9J08. The source data are provided with this paper.

## Abbreviations

AXE: acetyl xylan esterase
FAE: ferulic acid esterase
ESTHER: ESTerases and alpha/beta-Hydrolase Enzymes and Relatives
CE: carbohydrate esterase
CAZy: Carbohydrate-Active enZymes
*Al*AXEA: *A. luchuensis* AXE A
*Ao*AXEB: *A. oryzae* AXE B
*Ao*AXEC: *A. oryzae* AXE C
*Tf*FAEA: *T. funiculosus* FAE A
*Ao*FaeD: *A. oryzae* FAE D
*Ao*AXEA: *A. oryzae* AXE A
Endo-H: endo-β-*N*-acetylglucosaminidase H
*Tl*Xyn: *T. lanuginosus* xylanase
RMSD: root mean square deviations
*Lt*PHBase: *L. thermophila* poly[(*R*)-3-hydroxybutyric acid] depolymerase
*Aa*FaeD: *A. alcalophilum* FAE D
*As*FaeE: *A. sydowii* FAE E
*Tm*EstA: *T. maritima* thermophilic esterase
2AcX_3_: 2-acetylated xylotriose
3AcX_3_: 3-acetylated xylotriose

## Notes

### Competing Interest Statement

The authors have declared no competing interest.

## References

1 Scheller HV & Ulvskov P (2010) Hemicelluloses. Annu Rev Plant Biol 61, 263–289.

2 Biely P (2012) Microbial carbohydrate esterases deacetylating plant polysaccharides. Biotechnol Adv 30, 1575–1588.

3 Biely P, Westereng B, Puchart V, de Maayer P & Cowan DA (2014) Recent Progress in Understanding the Mode of Action of Acetylxylan Esterases. J Appl Glycosci (1999) 61, 35–44.

4 Puchart V & Biely P (2023) Microbial xylanolytic carbohydrate esterases. Essays Biochem 67, 479–491.

5 Nardini M & Dijkstra BW (1999) α/β Hydrolase fold enzymes: the family keeps growing. Curr Opin Struct Biol 9, 732–737.

6 Chatonnet A, Perochon M, Velluet E & Marchot P (2023) The ESTHER database on alpha/beta hydrolase fold proteins - An overview of recent developments. Chem Biol Interact 383, 110671.

7 Drula E, Garron ML, Dogan S, Lombard V, Henrissat B & Terrapon N (2022) The carbohydrate-active enzyme database: functions and literature. Nucleic Acids Res 50, D571–D577.

8 Adesioye FA, Makhalanyane TP, Biely P & Cowan DA (2016) Phylogeny, classification and metagenomic bioprospecting of microbial acetyl xylan esterases. Enzyme Microb Technol 93–94, 79–91.

9 Li X, Griffin K, Langeveld S, Frommhagen M, Underlin EN, Kabel MA, de Vries RP & Dilokpimol A (2020) Functional validation of two fungal subfamilies in carbohydrate esterase family 1 by biochemical characterization of esterases from uncharacterized branches. Front Bioeng Biotechnol 8, 542687.

10 Chung HJ, Park SM, Kim HR, Yang MS & Kim DH (2002) Cloning the gene encoding acetyl xylan esterase from *Aspergillus ficuum* and its expression in *Pichia pastoris*. Enzyme Microb Technol 31, 384–391.

11 Koseki T, Furuse S, Iwano K, Sakai H & Matsuzawa H (1997) An *Aspergillus awamori* acetylesterase: purification of the enzyme, and cloning and sequencing of the gene. Biochemical Journal 326, 485–490.

12 Koseki T, Miwa Y, Akao T, Akita O & Hashizume K (2006) An *Aspergillus oryzae* acetyl xylan esterase: Molecular cloning and characteristics of recombinant enzyme expressed in *Pichia pastoris*. J Biotechnol 121, 381–389.

13 Bauer S, Vasu P, Persson S, Mort AJ & Somerville CR (2006) Development and application of a suite of polysaccharide-degrading enzymes for analyzing plant cell walls. Proc Natl Acad Sci U S A 103, 11417–11422.

14 Puchart V, Agger JW, Berrin JG, Várnai A, Westereng B & Biely P (2016) Comparison of fungal carbohydrate esterases of family CE16 on artificial and natural substrates. J Biotechnol 233, 228–236.

15 Kato T, Shiono Y & Koseki T (2021) Identification and characterization of an acetyl xylan esterase from *Aspergillus oryzae*. J Biosci Bioeng 132, 337–342.

16 Zhang Y, Ding HT, Jiang WX, Zhang X, Cao HY, Wang JP, Li CY, Huang F, Zhang XY, Chen XL, Zhang YZ & Li PY (2021) Active site architecture of an acetyl xylan esterase indicates a novel cold adaptation strategy. Journal of Biological Chemistry 297, 100841.

17 Akoh CC, Lee GC, Liaw YC, Huang TH & Shaw JF (2004) GDSL family of serine esterases/lipases. Prog Lipid Res 43, 534–552.

18 Komiya D, Hori A, Ishida T, Igarashi K, Samejima M, Koseki T & Fushinobu S (2017) Crystal structure and substrate specificity modification of acetyl xylan esterase from *Aspergillus luchuensis*. Appl Environ Microbiol 83.

19 Koseki T, Handa H, Watanabe Y ya, Ohtsuka M & Shiono Y (2016) An unusual feruloyl esterase from *Aspergillus oryzae*: two tryptophan residues play a crucial role for the activity. J Mol Catal B Enzym 133, S560–S568.

20 Krissinel E & Henrick K (2007) Inference of macromolecular assemblies from crystalline state. J Mol Biol 372, 774–797.

21 Thomas GM, Quirk S, Huard DJE & Lieberman RL (2022) Bioplastic degradation by a polyhydroxybutyrate depolymerase from a thermophilic soil bacterium. Protein Science 31, e4470.

22 Phienluphon A, Kondo K, Mikami B, Teo KSK, Saito K, Watanabe T, Nagata T & Katahira M (2024) Structure-Based Characterization and Improvement of an Enzymatic Activity of Acremonium alcalophilum Feruloyl Esterase. ACS Sustain Chem Eng 12, 3831–3840.

23 Phienluphon A, Kondo K, Mikami B, Nagata T & Katahira M (2023) Structural insights into the molecular mechanisms of substrate recognition and hydrolysis by feruloyl esterase from Aspergillus sydowii. Int J Biol Macromol 253, 127188.

24 Levisson M, Sun L, Hendriks S, Swinkels P, Akveld T, Bultema JB, Barendregt A, van den Heuvel RHH, Dijkstra BW, van der Oost J & Kengen SWM (2009) Crystal structure and biochemical properties of a novel thermostable esterase containing an immunoglobulin-like domain. J Mol Biol 385, 949–962.

25 Berg JM, Gatto GJ, Hines J, Tymoczko JL & Stryer L (2023) 6. Enzyme Catalytic Strategies. In Biochemistry 10th Edition, pp. 179–209. W. H. Freeman, San Francisco.

26 Ichishima E (2016) Development of enzyme technology for *Aspergillus oryzae*, *A. sojae*, and *luchuensis*, the national microorganisms of Japan. Biosci Biotechnol Biochem 80, 1681– 1692.

27 Faulds CB & Williamson G (1994) Purification and characterization of a ferulic acid esterase (FAE-III) from *Aspergillus niger*: Specificity for the phenolic moiety and binding to microcrystalline cellulose. Microbiology (N Y) 140, 779–787.

28 Kabsch W (2010) XDS. Acta Crystallogr Sect D-Struct Biol 66, 125–132.

29 Evans PR & Murshudov GN (2013) How good are my data and what is the resolution? Acta Crystallogr Sect D-Struct Biol 69, 1204–1214.

30 Vagin A & Lebedev A (2015) MoRDa, an automatic molecular replacement pipeline. Acta Crystallogr A Found Adv 71, s19–s19.

31 Liebschner D, Afonine P V., Baker ML, Bunkoczi G, Chen VB, Croll TI, Hintze B, Hung LW, Jain S, McCoy AJ, Moriarty NW, Oeffner RD, Poon BK, Prisant MG, Read RJ, Richardson JS, Richardson DC, Sammito MD, Sobolev O V., Stockwell DH, Terwilliger TC, Urzhumtsev AG, Videau LL, Williams CJ & Adams PD (2019) Macromolecular structure determination using X-rays, neutrons and electrons: recent developments in Phenix. Acta Crystallogr Sect D-Struct Biol 75, 861–877.

32 Cowtan K (2006) The Buccaneer software for automated model building. 1. Tracing protein chains. Acta Crystallogr Sect D-Biol Crystallogr 62, 1002–1011.

33 Murshudov GN, Skubák P, Lebedev AA, Pannu NS, Steiner RA, Nicholls RA, Winn MD, Long F & Vagin AA (2011) REFMAC5 for the refinement of macromolecular crystal structures. Acta Crystallogr D Biol Crystallogr 67, 355–367.

34 Liebschner D, Afonine P V., Moriarty NW, Poon BK, Sobolev O V., Terwilliger TC & Adams PD (2017) Polder maps: Improving OMIT maps by excluding bulk solvent. Acta Crystallogr D Struct Biol 73, 148–157.

35 Tamura K, Stecher G & Kumar S (2021) MEGA11: Molecular evolutionary genetics analysis version 11. Mol Biol Evol 38, 3022–3027.

36 Edgar RC (2004) MUSCLE: Multiple sequence alignment with high accuracy and high throughput. Nucleic Acids Res 32, 1792–1797.

37 Eberhardt J, Santos-Martins D, Tillack AF & Forli S (2021) AutoDock Vina 1.2.0: New docking methods, expanded force field, and Python bindings. J Chem Inf Model 61, 3891–3898.

38 Okano H, Hong X, Kanaya E, Angkawidjaja C & Kanaya S (2015) Structural and biochemical characterization of a metagenome-derived esterase with a long N-terminal extension. Protein Science 24, 93–104.

39 Kroon PA, Williamson G, Fish NM, Archer DB & Belshaw NJ (2000) A modular esterase from *Penicillium funiculosum* which releases ferulic acid from plant cell walls and binds crystalline cellulose contains a carbohydrate binding module. Eur J Biochem 267, 6740– 6752.

40 Crepin VF, Faulds CB & Connerton IF (2003) A non-modular type B feruloyl esterase from *Neurospora crassa* exhibits concentration-dependent substrate inhibition. Biochemical Journal 370, 417–427.

41 Gordillo F, Caputo V, Peirano A, Chavez R, Van Beeumen J, Vandenberghe I, Claeyssens M, Bull P, Ravanal MC & Eyzaguirre J (2006) *Penicillium purpurogenum* produces a family 1 acetyl xylan esterase containing a carbohydrate-binding module: characterization of the protein and its gene. Mycol Res 110, 1129–1139.

