## Supplementary Tables and Figures for "Identification and structural characterization of a novel acetyl xylan esterase from *Aspergillus oryzae*"

**Supplementary Table S1. Results of *N*-glycosylation site prediction.**

| Site | Sequence | Potential | Jury agreement | N-Glyc result |
|---|---|---|---|---|
| N42 | NFTL | 0.6271 | 8/9 | + |
| N248 | NTTK | 0.7838 | 9/9 | +++ |
| N295 | NGTY | 0.4827 | 5/9 | - |
| N321 | NQSE | 0.4359 | 6/9 | - |

Predicted by the NetNGlyc 1.0 server (https://services.healthtech.dtu.dk/services/NetNGlyc-1.0/) [1].

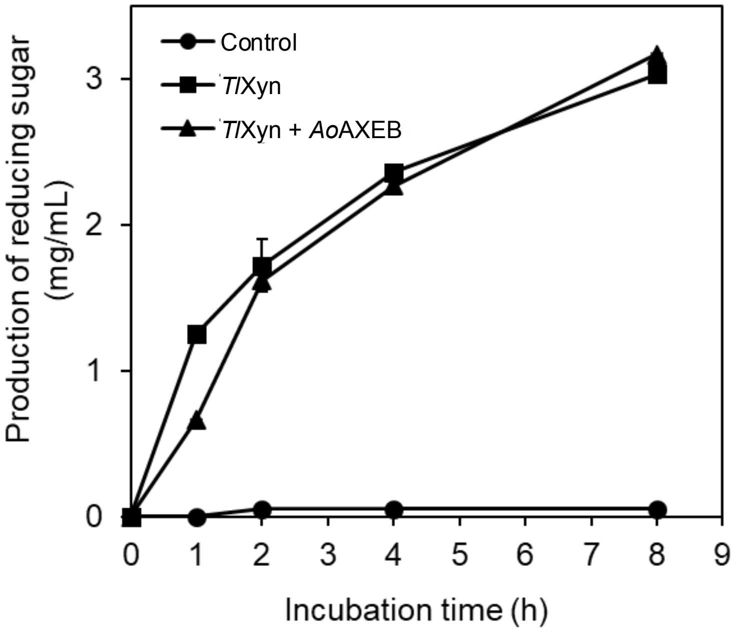


**Supplementary Fig. S1.** Effect of *Ao*AXEB on the production of reducing sugar derived from arabinoxylan degradation by *T. lanuginosus* xylanase (*Tl*Xyn). Symbols: closed circles, nontreated arabinoxylan; closed squares, treated arabinoxylan with *Tl*Xyn; closed triangles, treated arabinoxylan with *Tl*Xyn plus *Ao*AXEB.

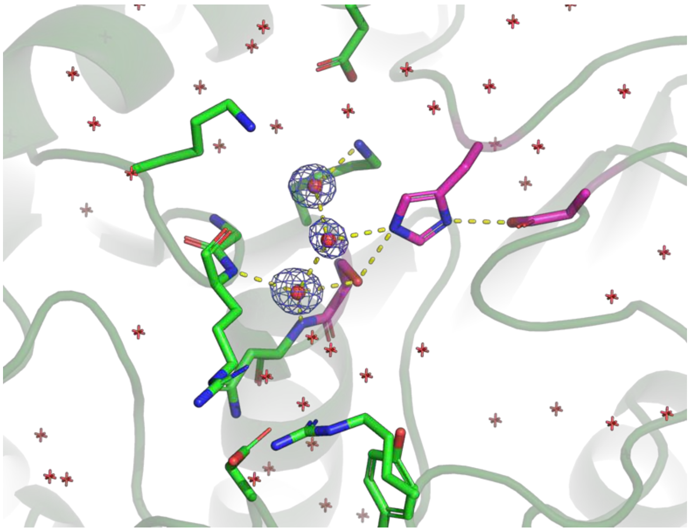

**Supplementary Fig. S2.** Active site of the apo structure. A Polder map (5σ) for three water molecules near the catalytic center. Other water molecules are shown as red crosshairs. The catalytic triad and other residues in the active site are shown as magenta and green sticks, respectively.

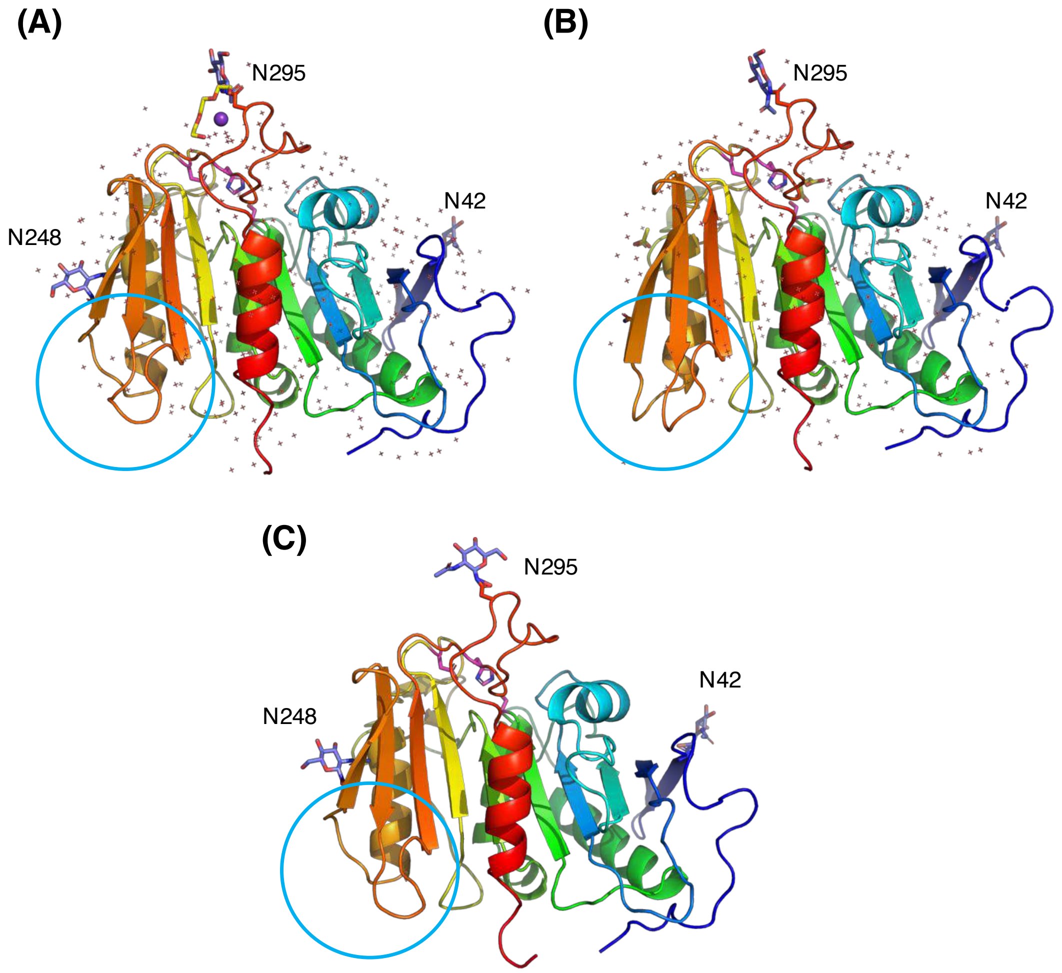


**Supplementary Fig. S3.** Comparison of the crystal structures of *Ao*AXEB with predicted structure. (A) Apo structure. Polyethylene glycol molecule (triethylene glycol) and a sodium ion bound on the protein surface are shown as yellow sticks and a purple sphere, respectively. (B) Succinate complex structure. Bound succinate and acetate molecules are shown as yellow sticks. (C) A protein structure (model_0) predicted using the AlphaFold server (beta) (https://alphafoldserver.com) [2] on 1 August, 2024. Mature protein sequence of *Ao*AXEB (residues 19–326) with *N*-glycosylation of a single *N*-acetylglucosamine at N42, N248, and N295 was used as a query. The catalytic triad residues are shown as magenta sticks, and water molecules in the crystal structures are shown as red crosshairs. Areas where differences were observed in these structures are circled with cyan.

**Supplementary References**

1 Gupta R & Brunak S (2002) Prediction of glycosylation across the human proteome and the correlation to protein function. *Pac Symp Biocomput* **7**, 310–322.

2 Abramson J, Adler J, Dunger J, Evans R, Green T, Pritzel A, Ronneberger O, Willmore L, Ballard AJ, Bambrick J, Bodenstein SW, Evans DA, Hung CC, O'Neill M, Reiman D, Tunyasuvunakool K, Wu Z, Žemgulytė A, Arvaniti E, Beattie C, Bertolli O, Bridgland A, Cherepanov A, Congreve M, Cowen-Rivers AI, Cowie A, Figurnov M, Fuchs FB, Gladman H, Jain R, Khan YA, Low CMR, Perlin K, Potapenko A, Savy P, Singh S, Stecula A, Thillaisundaram A, Tong C, Yakneen S, Zhong ED, Zielinski M, Žídek A, Bapst V, Kohli P, Jaderberg M, Hassabis D & Jumper JM (2024) Accurate structure prediction of biomolecular interactions with AlphaFold 3. *Nature 2024 630:8016* **630**, 493–500.
